# Severe population collapses and species extinctions in multi-host epidemic dynamics

**DOI:** 10.1101/128363

**Authors:** Sergei Maslov, Kim Sneppen

**Affiliations:** Department of Bioengineering and Carl R. Woese Institute for Genomic Biology, University of Illinois at Urbana-Champaign, Urbana, IL 61801, USA; Center for Models of Life, Niels Bohr Institute, University of Copenhagen, 2100 Copenhagen, Denmark

## Abstract

Most infectious diseases including more than half of known human pathogens are not restricted to just one host, yet much of the mathematical modeling of infections has been limited to a single species. We investigate consequences of a single epidemic propagating in multiple species and compare and contrast it with the endemic steady state of the disease. We use the two-species Susceptible-Infected-Recovered (SIR) model to calculate the severity of post-epidemic collapses in populations of two host species as a function of their initial population sizes, the times individuals remain infectious, and the matrix of infection rates. We derive the criteria for a very large, extinction-level, population collapse in one or both of the species. The main conclusion of our study is that a single epidemic could drive a species with high mortality rate to local or even global extinction provided that it is co-infected with an abundant species. Such collapse-driven extinctions depend on factors different than those in the endemic steady state of the disease.

## INTRODUCTION

Models of pathogen dynamics for the most part include only a single host species [1] in spite of the fact that pathogens typically infect multiple species. For example, more than half of human pathogens are known to be shared with at least one animal species [2, 3]. Famous examples of diseases with multiple host species include cholera (*Vibrio cholerae*) commensal in a number zooplankton species [4, 5], bubonic plague (Yersinia pestis) co-infecting and spreading between humans and rats [6], and more recently the avian influenza virus [7]. The steady state of dynamical equations where multiple hosts are infected by the same pathogen was previously considered by Dobson [8]. This important study addressed the interplay between the diversity of hosts and the stability of disease’s endemic state. In our study we chose to focus on the transient (as opposed to the steady state) dynamics of a single epidemic as it is spreading in several species. While our mathematical formalism can be easily generalized to an arbitrary number of species, our main results can be already demonstrated for just two species. In what follows we use only this simpler two-species model.

The dynamic of a single epidemic of a disease is often described in terms of the SIR (Susceptible – Infected – Removed) model [11] and its variants: *dS/dt* = -*βS ˙ I*, *dI/dt* = *βS ˙ I - γI* and *dD/dt* = *γI*. In this model individual members of the population susceptible to disease (*S*) become infected (*I*), and are subsequently removed from the pool spreading the disease due to either their death (*D*) or newly acquired immunity. While from the mathematical perspective there is no difference between death and complete resistance to disease, only the former results in population collapses that are the focus of our study. A well known property of the single species SIR model [11] is that in the course of the first epidemic the population of the host species drops to a much lower level than its ultimate steady state population in the endemic state of the diseases. When this population collapse is especially severe the host species is vulnerable to either local or even complete extinction. Such collapses along with extinction events triggered by them are the main focus of our study.

The mass-action equations describing the dynamics of transitions between the three states of the SIR model can be described by a single key parameter, *R*_0_ (equal to *βS*(0)*/γ* in the notation used above), called the basic reproduction number or the epidemiological threshold. It is defined as the number of new infections caused by each infected individual at the very start of the epidemic when the density of susceptible individuals is still close to *S*(0) - its value at the start of the epidemic. Thus for *R*_0_ *>* 1 the infection started by a very small number of infected individuals will (at least initially) exponentially amplify and ultimately reduce the size of the susceptible population. In the opposite case *R*_0_ *<* 1 the initial infection will quickly fizzle out and the population size will remain virtually unchanged. As the epidemic spreads, the number of susceptible targets declines, ultimately leaving *S*(collapse) survivors. For *R*_0_ *≫* 1 the population collapse is given by the exponential function of *R*_0_: ≃ *S*(collapse) *S*(0) exp[-*R*_0_]) for 0 *< I*(0) ≫ *S*(0). The exponential decline in the number of these survivors of an epidemic as a function of *R*_0_ can be derived through eliminating the non-linear term in the SIR model by measuring time in units of the number of deaths: *dS/dD* = – (*β/γ*)*S*. Thus *S*(*t*) = *S*(0) exp[-*βD*(*t*)*/γ*] leading to the final number of survivors after the epidemic ran its course *S*(collapse) = *S*(0) exp[-*βD*(collapse)*/γ*] = *S*(0) exp[-*β*(*S*(0)-*S*(collapse))*/γ*] *≃ S*(0) exp[-*βS*(0)*/γ*] = *S*(0) exp[-*R*_0_].

It is illustrative to compare the size of the population after a collapse with its steady state value in the endemic state of the disease. Such endemic state requires a constant source of susceptible individuals which is traditionally realized by adding a small birth term to the SIR model (see e.g. Ref. [8]). The collapse *S*(0) *→ S*(0)*/γ*] = *S*(0) exp[– *R*_0_] dramatically overshoots the endemic steady state population density *S*(steady state) = *S*(0)*/R*_0_ In the endemic state of the disease each infected individual transmits it to exactly one other susceptible individual thereby keeping a permanent infection going without exponential expansion or decay. Hence, 1 = *S*(steady state)*β/γ* = *S*(steady state)*R*_0_*/S*(0).

A classic example of a pathogen-host ecosystem over-shooting its steady state immediately after the first epidemic can be found e.g. in the experiments carried out in Ref. [12]: when a new phage was introduced into a bacterial population dominated by susceptible strains resulted in a bacterial population drop by roughly 5 orders of magnitude followed by a slow recovery to the steady state which is only one order of magnitude lower than the population at the start of the experiment. Similar contrast between the initial population collapse is possible for epidemics of airborne diseases such as measles or small pox where *R*_0_ could exceed 10 in an immunological naive population. While measles or small pox do not always kill infected individuals, if a similarly contagious disease that is 100% lethal to its hosts was to emerge, the initial epidemic-induced collapse exp(–*R*_0_) = exp(–10) *∼* 5 ˙ 10^−*5*^ would reduce host’s population to much below its long-term steady state level of 1*/R*_0_ *∼* 1*/*10 achieved when (or if) such disease would become endemic. One expects a local extinction of the species if the population of survivors after the epidemic, *S*(collapse) ≃ *S*(0) exp(–*R*_0_), drops below one individual.

In this paper we model a single epidemic of a disease infecting multiple host species and investigate how its transient dynamics can result in a severe collapse or even local extinction of either of these species. Such a scenario is realistic because epidemics routinely spill over to other species, that is to say, diseases transiently or permanently transverse species boundaries. For example, several Ebola epidemics in wild gorilla groups in central Africa happened between 2002 and 2003 resulted in 90%-95% reduction in gorilla populations [13]. Such local near-extinction collapses have been blamed on ongoing spillover of the Ebola virus from its reservoir host, fruit eating bats, subsequently amplified by ape-to-ape virus transmission [14].

## METHODS

The two-host SIR model describes the disease propagation in species 1 and 2 via the following system of ODEs:

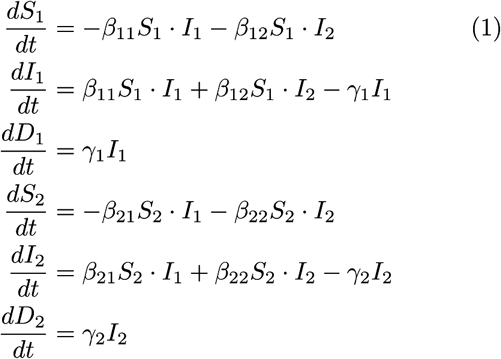

Here we assume density-dependent transmission mechanism characteristic of non-sexually or vector-transmitted diseases in well-mixed populations. We use the traditional notation [8] where *β*_*ij*_ is the matrix of transmission rates from species *j* to species *i*.

The SIR equations remain the same if instead of dying infected individuals recover with full immunity. However, since the focus of our study is on population collapses we choose to interpret *γ*_*i*_ as the death rate of infected individuals of the species *i* (see Discussion for a more general case including both death and recovery). *S*_*i*_, *I*_*i*_ and *D*_*i*_ refer to population densities of susceptible, infected, and dead individuals in each of two species correspondingly. Our model describes the time course of a single epidemic during which we ignore births of new susceptible individuals. This approximation is justified when the time from infection to death is fast compared to other timescales in the system.

The epidemic is initiated at time *t* = 0 with a very small numbers of infected hosts in either one or both species: *I*_1_(0) *≫ S*_1_(0), and *I*_2_(0) *≫ S*_2_(0). In this limit the resulting population dynamics is independent of the exact values of *I*_1_(0) and *I*_2_(0).

## RESULTS

We numerically simulated the time dynamics of Eqs. 1, see Fig. 1. To compare the results of a single epidemic to the endemic state of the disease we added a small birth term with saturation given by 0.01 ˙ *S*_*i*_(*t*) ˙ (1 - *S*_*i*_(*t*)) to the right hand side of the equations for *dS*_*i*_(*t*)*/dt*. We also added even smaller natural (non-disease related) death term -0.0001 ˙ *S*_*i*_(*t*) to equations for *dS*_*i*_(*t*)*/dt* and a similar death term - 0.0001 ˙ *I*_*i*_(*t*) to equations for *dI*_*i*_(*t*)*/dt*. This term ensures the flow of newly born susceptible individuals without affecting much either the population collapse after the first epidemic nor the long-term steady state of the system. We then start our simulations at a pre-epidemic susceptible population *S*_*i*_(0) = 0.99 *∼* 1. The general steady state analysis of these equations has been carried out by Dobson [8]. The birth term used in our study differs slightly from that used in Ref. [8] as we assume that infected individuals are infertile. When the growth rate is small (i.e. ≫ *γ, ≪*) these choices do not significantly influence the collapse ratio (data not shown). In our interpretation of the SIR model the “removed” individuals are dead and thus (naturally) not included in the birth term. This would change if infected individuals recover with a full immunity and are capable of giving birth [8].

**FIG. 1.**
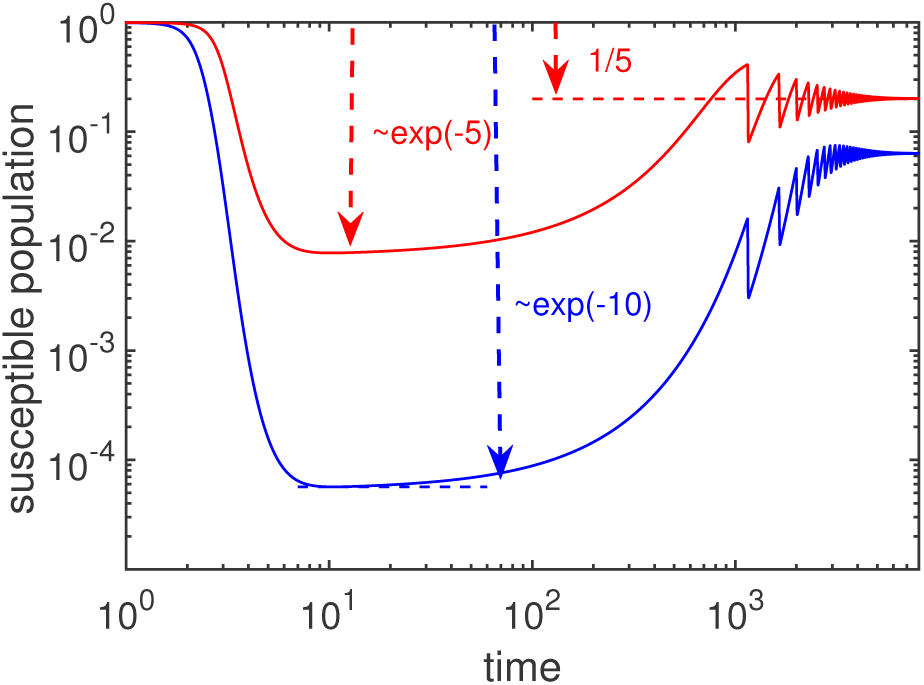
Simulation of Eqs. 1 for populations of two species *S*_1_(*t*) (blue) and *S*_2_(*t*) (red) susceptible to infection by a single pathogen with (cross)infection rates *β*_11_ = *β*_12_ = *β*_22_ = 5, *β*_21_ = 0, and *γ*_1_ = *γ*_2_ = 1. At the start of the simulation *S*_1_(0) = *S*_2_(0) = 1 and *I*_1_(0) = 0, *I*_2_(0) = 10^−6^. In order for equations to have an endemic steady state we added small birth and natural death terms as described in the text. Right and left red arrows respectively point to the predicted steady state of the second population *S*_2_(steady state) = 1*/*5, and its population immediately after the first epidemic *S*_2_(collapse) ≃ exp(–5). The blue arrow highlights a much more severe post-epidemic collapse of the first species: *S*_1_(collapse) *≃* exp(-10).

We first consider a simple scenario when the transmission is unidirectional 2 *→* 1. In this case an epidemic started in the species 2 would spill over to the species 1 and cause its population to collapse but not vice versa. A case study is presented in Fig. 1 where we plot time-courses of susceptible populations *S*_1_(*t*) (blue) and *S*_2_(*t*) (red) defined by equations 1 with *β*_11_ = *β*_12_ = *β*_22_ = 5, *β*_21_ = 0, and *γ*_1_ = *γ*_2_ = 1.

For the case explored in Fig. 1 the population dynamics of the second species is independent of the first one. Thus, like in a single species case outlined before, its epidemics is characterized by the basic reproduction number *S*(0)*β*_22_*/γ*_2_ = 5. In the endemic steady state its population is expected to be close to *S*_2_(steady state) = 1*/*5 (the red dashed line in Fig. 1), while its initial post-epidemic collapse population to be approximately equal to *S*_2_(collapse) = exp(–5) (the red arrow on the left of Fig. 1). Noticeably the species 1 (shown with blue) positioned “downstream” of the epidemics in the species 2, is exposed to a much worse disease outbreak than the species 2. Its population collapses down to *S*_1_(collapse) = exp(–10) *≫ S*_2_(collapse). This amplification of outbreaks in two-or multi-host epidemics can be described by the general theoretical framework described below.

The equations 1 include both the unidirectional case discussed above, and the possibility that there is cross-infections in both directions. The equations can be solved by introducing two “composite death toll” variables 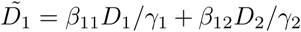 and 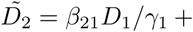. When these variables are used instead of time for each of two species, their susceptible populations follow a simple exponential decay 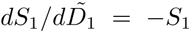 and *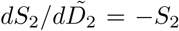* ending at their new post-collapse densities given by 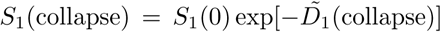 and *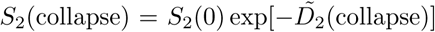*. We quantify the impact of the epidemic on populations of each of two species by Γ_*i*_ defined by exp(–Γ_*i*_) = *S*_1_(collapse)*/S*_1_(0).

Since at the end of the epidemic the number of infected individuals is equal to zero, the overall death tolls are given by *D*_1_ (collapse) = *S*_1_(0)-*S*_1_ (collapse) and *D*_2_(collapse) = *S*_2_(0) –*S*_2_(collapse). The fractions of two populations that died during the epidemic *ρ*_1_ = *D*_1_(collapse)*/S*_1_(0) = 1 –*S*_1_(collapse)*/S*_1_(0) = 1 – exp(–Γ_1_) and *ρ*_2_ = *D*_2_(collapse)*/S*_2_(0) = 1 – *S*_2_(collapse)*/S*_2_(0) = 1– exp(−Γ_2_) are then selfconsistently determined by

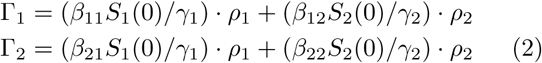

This non-linear system of equations can be numerically (e.g. iteratively) solved for *ρ*_*i*_ = 1 − exp(-Γ_1_). The solution is fully determined by the *collapse matrix*:

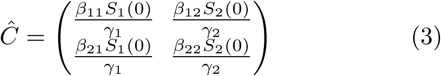

Note, that this collapse matrix, describing the cumulative *aftermath of an epidemic* is subtly yet critically different from the commonly used “next generation matrix” [9, 10] describing the dynamics at the very *start of the epidemic*:

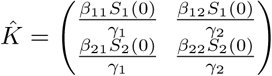

One can show that a non-zero collapse with Γ _*i*_ *>* 0 in any of the species is possible if and only if the largest eigen-value of the matrix 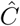 exceeds 1. This does not contradict the classic result [8–10] that the basic reproduction number of the epidemic, *R*_0_ *>* 1, is equal to the largest eigen-value of the next generation matrix 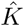. The agreement is ensured by the mathematical fact that the collapse and the next generation matrices are connected to each other by the similarity transformation 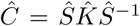 and thus have identical eigenvalues. Here 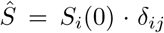 is the diagonal matrix of initial species abundances.

In the limit where population collapses in both species are large (*ρ*_1_ ∼ 1 and *ρ*_2_ ∼ 1), the Eqs. 2 predict the logarithm of collapse ratios in each of two populations to be given by a simple sum of matrix elements of the collapse matrix: Γ_1_ = *C*_11_ + *C*_12_ and Γ_2_ = *C*_21_ + *C*_22_. In other words, the overall fraction of survivors exp(–Γ_*i*_) is given by a product of survival probabilities in infections transmitted by the members of its own species and those of the opposite species. This is illustrated by the case of unidirectional transmission shown in Fig. 1, where the “downstream” species 1 collapses by a factor exp(–10) = exp(–5) ˙ *exp*(–5) = exp(–*C*_11_) ˙ exp(–*C*_12_), while the “upstream” species 2 collapses only by a factor exp(–5) = exp(– *C*_22_). If the disease was able to spread equally in both directions, both species would suffer equally large collapses *∼* exp(-10).

Figs. 2-3 show the decimal logarithm (as opposed to the natural one) of the species 1 collapse ratio log_10_(*S*_1_(0)*/S*_1_(collapse)) = − Γ_1_*/* ln(10) for different combinations of parameters. In Fig. 2 we examine the logarithmic magnitude of the species 1 collapse, as a function of initial susceptible population sizes of both species (panel (a)) and (cross)infections rates (panel (b)). Panel (a) plots log_10_(*S*_1_(0)*/S*_1_(collapse)) as a function of the initial populations *S*_1_(0) and *S*_2_(0) in a system where *γ*_1_ = *γ*_2_ = 1 and *β*_11_ = *β*_12_ = 0.3, and *β*_22_ = *β*_21_ = 3. White line is the predicted epidemic threshold below which the largest eigenvalue of the collapse matrix Ĉ falls below 1. White dot marks the population sizes *S*_1_(0) = 1 and *S*_2_(0) = 10 used in the panel (b), which shows log_10_(*S*_1_(0)*/S*_1_(collapse)) at these population sizes and variable infection rates *β*_11_ = *β*_12_, and *β*_22_ = *β*_21_. Note that Figs. 2 and 3 shows the decimal logarithm of the collapse ratio. Thus for a population of, for example, 10_5_ individuals, a collapse value greater than 5 (yellow-to-red colors in our Figures 2 and 3) indicates a likely local extinction threshold for species 1 defined by the epidemic reducing the population to (on average) *<* 1 surviving individuals.

**FIG. 2.**
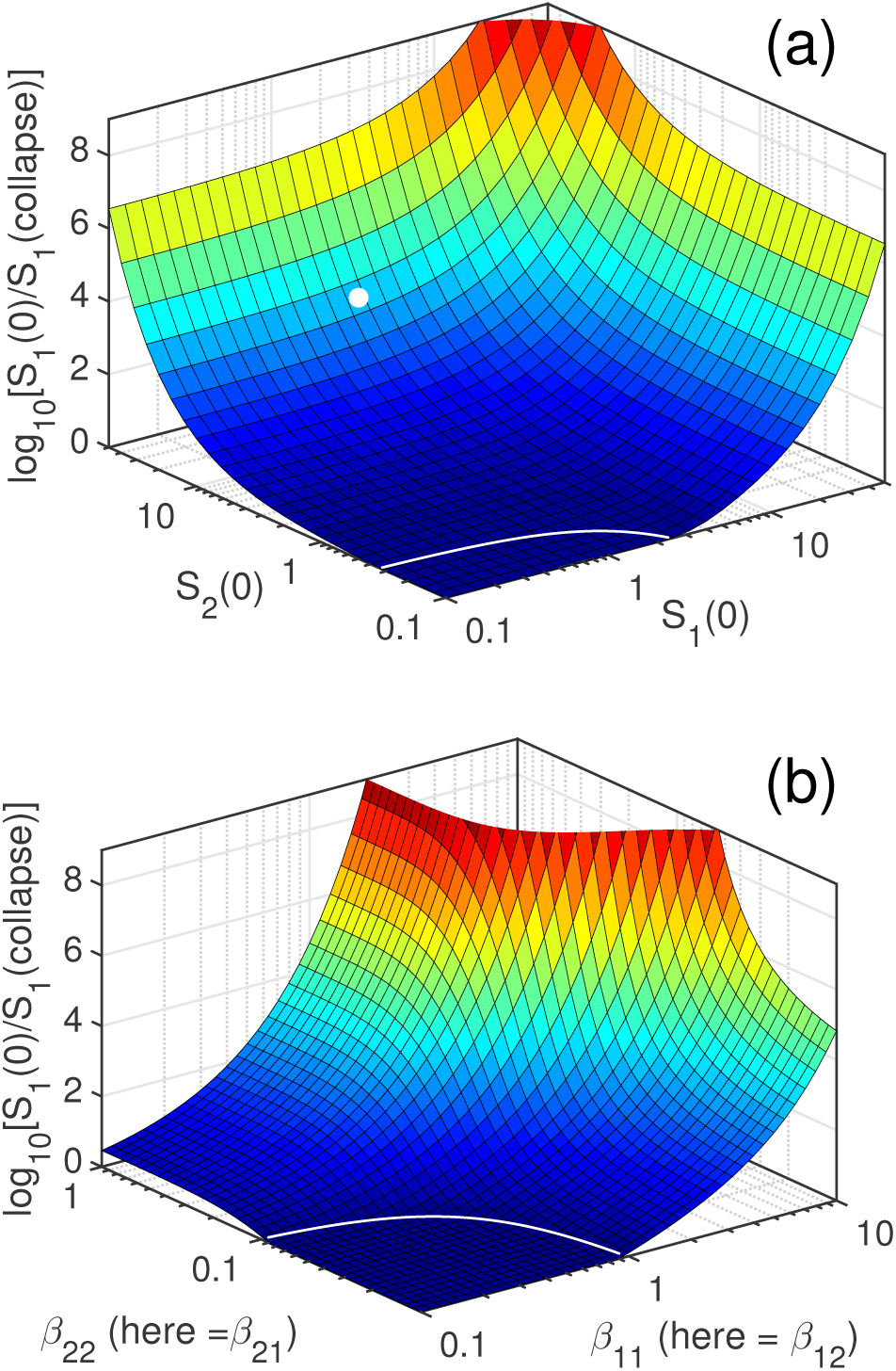
Decimal logarithm of the collapse ratio log_10_ (*S*_*1*_(0)*/S* (collapse)) = Γ_1_ */* ln(10) in the population 1 as a function of the two species population sizes (panel (a)) and infections rates (panel (b)). Panel (a) shows the collapse ratio as a function of the initial populations *S*_1_(0) and *S*_2_(0) in a system where *γ*_1_ = *γ*_2_ = 1 and *β*_11_ = *β*_12_ = 0.3, and *β*_22_ = *β*_21_ = 3. White line is the predicted epidemic thresh-old at which the largest eigenvalue of the collapse matrix *Ĉ* is equal to 1. Yellow-to-red colors indicate likely extinction of the species 1 with the initial population of 10^5^ susceptible individuals. White dot marks initial population sizes *S*_1_(0) = 1 and *S*_2_(0) = 10 used in panel (b), which shows the decimal logarithm of the collapse ratio calculated for these initial population sizes and variable infection rates *β*_11_ = *β*_12_, and *β*_22_ = *β*_21_. White line marks the predicted epidemic thresh-old.

In general, two host species in our model are characterized by different infection parameters and potentially highly asymmetric transmission rates. For example, for Ebola virus in bats and gorillas mentioned above [13, 14] cross infections are believed to be mediated primarily by bats’ droppings landing on fruits that gorillas eat. Thus the spread of the virus is generally uni-directional from species 2 (bats) to species 1 (gorillas). In Fig. 1 we simulated our model with *β*_21_ = 0. In Fig. 2 we examine how the magnitude of the post-epidemic drop in population sizes depends on parameters. Fig. 2a shows the dependence of the logarithmic collapse ratio in the species 1 (gorillas) on the size of the magnitude of cross-species collapse matrix element *C*_12_ = *β*_12_*S*_2_(0)*/γ*_2_ and the size of intra-species collapse number *C*_22_ within the species 2 (bat) population. To further illustrate our point we selected the basic collapse number in the population of gorillas to be well below the species 1 epidemic threshold if it was isolated from species 2 (*C*_11_ = 0.1 *<<* 1). Yet, we were anyway able to get an extinction-level collapse in the population of “gorillas” as long as the majority of bats were infected. It is important to note that our model equally well applies to the case where species 2 (bats) do not die in the course of the epidemic but are instead removed from the ranks of susceptible population by becoming immune to the disease (see the discussion for generalization of our mathematical formalism to in corporate recovery with immunity). The species 2 death rate *γ*_2_ in this case is simply the rate at which they acquire immunity and thus stop being infectious.

**FIG. 3.**
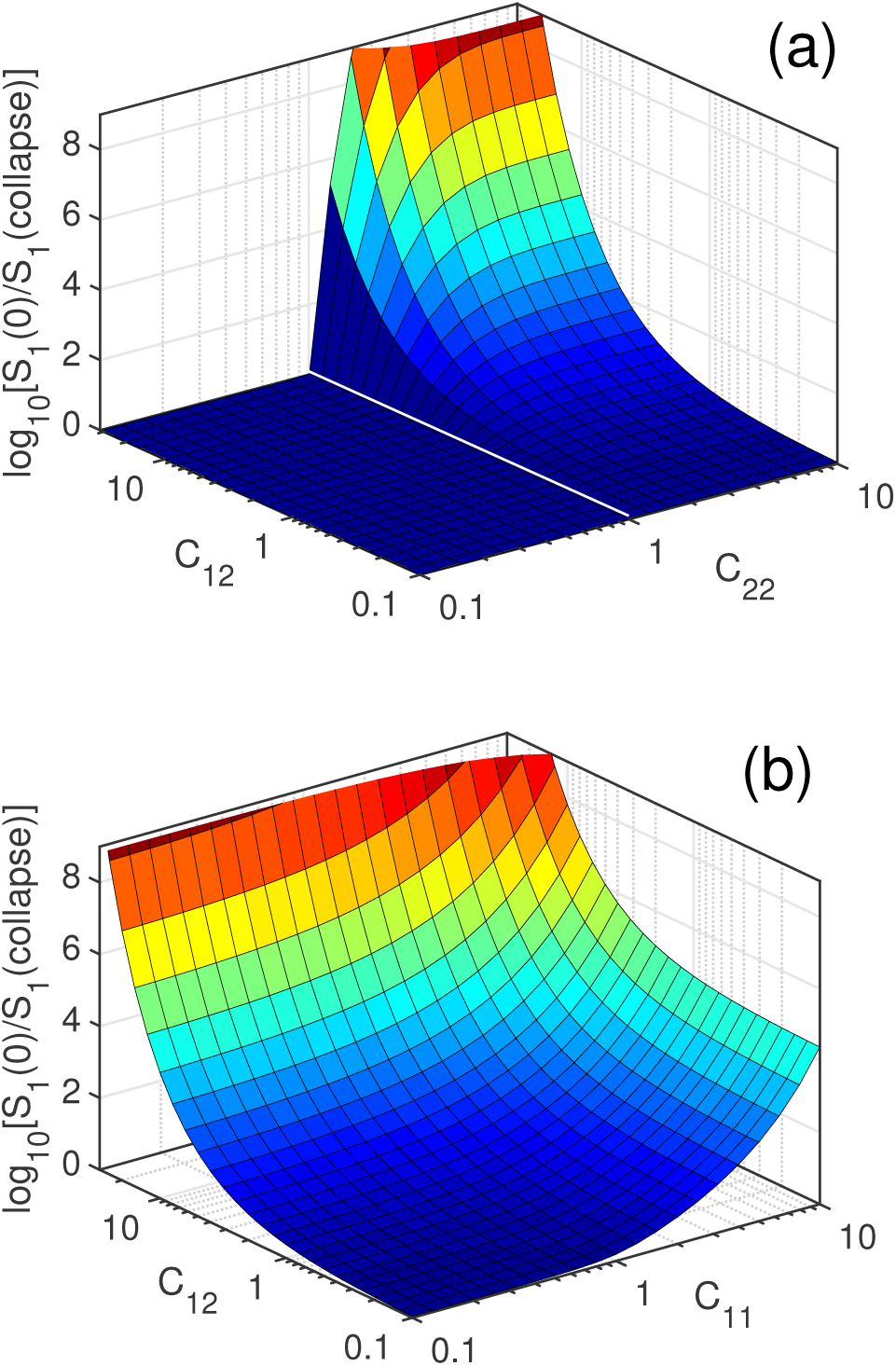
Collapse ratio *S*_1_(0)*/S*_1_(collapse of the population of the species 1 in the case of uni-directional transmission: *C*_21_ = 0, following an epidemic started with a very small number of infected individuals (*I*_1_(0) = *I*_2_(0) = 10^−6^). (a) Collapse ratio in the population of the species 1 with fixed intra-species collapse factor *C*_11_ = 0.1 as a function of the species 2 collapse number *C*_22_ and cross-species collapse number *C*_12_ quantifying disease transmission from species 2 to 1. White line is the predicted epidemic threshold below which the overall reproduction number falls below 1. (b) Collapse ratio the population of species 1, with a fixed value of *C*_22_ = 2 and variable *C*_11_ and *C*_12_. There is no epidemic threshold in this case as the basic reproduction number in the species 2 is selected to be larger than 1 so that the epidemic would always be able to start.

The properties of the system can be further analyzed in terms of a simple analytic expression obtained in the limit where Γ_1_ ≫ 1 and Γ_2_ ≫ 1 so that *ρ*_1_ *≃* 1 and *ρ*_2_ *≃* 1 (strictly speaking this is the limit of the model where Γ_*i*_, Γ_2_ *→ ∞*). In this case the Eq. 1 becomes simply

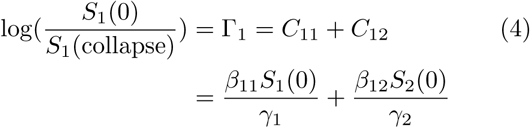

This limit approximately describes the simulations shown in Fig. 1 where the collapse of the first species is very close to *S*_1_(collapse) ≃ exp(–*C*_11_–*C*_22_) = exp(–10) (see blue arrow). Note that the population collapse in the species 1 described by the Eq. 4 does not depend on the impact of the epidemic on the species 2 population, corresponding to a near complete elimination of the susceptible population 2 (Γ_2_ ≫ 1). This limit can be seen in Fig. 2a as leveling off of the surviving fraction of the species 1 for large values *C*_22_ ≫ 1, corresponding to saturation of the reservoir of the species 2. Fig. 2b further explores this limit by plotting Γ_1_ as a function of *C*_11_ and *C*_12_ for a fixed “bat-to-bat” (within-species 2) collapse factor *C*_22_ = 2. In this case a large fraction of the population 2 (1– exp(–2) or 86%) becomes infected thus opening up plentiful opportunities (broad range of two other parameters of the model) for an extinction-level collapse of the population 1.

## DISCUSSION

Diseases are a real and constant danger for nearly any of the species on our planet, and are occasionally assumed to drive or facilitate extinction-scale events [18– 20]. This paper demonstrated that such events would be more likely when a lethal pathogen infects more than one host species. Above we explored a simple two species model subject to epidemic-driven population collapses and extinctions. The epidemic could be triggered by either the appearance of a new pathogen or a sudden increase in intra- or cross-species infection rates in a new ecological layout. As can be inferred from the Eq. 4 a severe population collapse of the species 1 is favored by an initially large population of the co-infecting species 2 (large *S*_2_(0)) that can stay infectious for a long time (*γ*_2_ small) resulting in a large cross-species collapse number *C*_12_.. Cross-species transmission could dramatically amplify the collapse due to within-species transmission which could even be characterized by a sub-critical value of *R*_0_(1 *→* 1) *<* 1 (*C*_11_ in our notation).

If a population would survive the first epidemic, one may speculate whether it would be sustainable in the long term endemic steady state. This was previously considered by [8], with the overall result was that coexistence of two or more species in the endemic steady state depends on multiple species-specific parameters. According to Ref. [8], the extinction of species in the endemic state is possible when intra-species transmission is high and it targets host species in the inverse order of their growth rates. That is to say, slowly growing species will go extinct first when they share pathogens with faster growing ones. Thus species survival in the endemic state of the disease depends on different parameters (growth rates) than in the initial epidemics (relative population sizes).

Our study suggests that transient epidemics of diseases provide species with powerful “weapons” against each other. Such weapons have been well documented in the microbial world where bacterial species co-infected by the same phage [15] fight ongoing battles with each other and their phage pathogen. Long history of such “red-queen” evolutionary dynamics can be inferred from many-layered defense and counter-defense mechanisms encoded within their genomes [16]. The use of shared diseases as a weapon have similarity to the apparent competition between multiple prey species sharing a common predator [17]. Our analysis extends these earlier results by including the impact of transient epidemics, and adding the possibility that permanently remove an otherwise fit predator.

An important example of cross-species interactions occurs when a single pathogen co-infects a highly abundant prey and its typically much its less abundant predator. Mapping the prey to species 2 in our model this situation would give rise to particularly large values of *C*_22_ and *C*_12_ which are both proportional to prey’s high population density *S*_2_(0). Our results suggest that such a disease may only need to be present during a relatively short period, in order to locally eliminate the less abundant predator species. Such disease would also result in a short-term population loss of the prey, but give it a long term gain in terms of eliminating the predator entirely. Alternatively, for the pathogen it would be evolutionary beneficial to be either completely benign or at least less deadly to its prey host, but much more lethal to its host’s predators. Indeed, by reducing predator population it increases prey (and hence its own) population. Thus in contrast to the classical single host results of [23, 24], our analysis suggests that it is not always beneficial for a disease to become more benign to *all of its hosts.*

Diseases often leave a substantial fraction of survivors, and their epidemics only cause a finite-size collapse in populations of their hosts. Somewhat counterintuitively this may increase the diversity of the host ecosystem by allowing hosts to bypass the competitive exclusion principle, according to which only the single fastest growing species survives in the long run. One example we investigated before [21] is the negative density-dependent selection in which phage epidemics preferentially spread in bacterial species or strains with large populations (so called “Kill-the-Winner” principle [22]) thereby leading to their abrupt and severe collapse.

Since the focus of this study is on extinction level population collapses, above we considered an extreme case of a disease with 100% mortality. Yet our results can be readily extended to a more general case in which a fixed fraction *x*_*i*_ of infected individuals of species *i* die, while 1–*x*_*i*_ - recover with full immunity. As was discussed above, for the purposes of the SIR mathematical model without birth these two outcomes are identical. Let *γ*_*i*_ denote the overall rate of death and recovery with immunity. Out of a fraction 1 – exp(-Γ_*i*_) removed from the corresponding susceptible population, *x*_*i*_(1 − exp(-Γ_*i*_)) actually died, while (1–*x*_*i*_)(1–exp(-Γ_*i*_)) survived. Thus by the end of the first epidemic the overall (both susceptible and immune) surviving population fraction is given by 1–*x*_*i*_ + *x*_*i*_ exp(–Γ_*i*_), where as before Γ_*i*_ is determined by the Eq. 2. Coming back to the bats and gorillas example considered above one can have a situation in which the Ebola virus is rather deadly (*x*_1_ ≃1) for one of the species (gorillas), while being mild in another (*x*_1_ ≃ 0) (bats [14]). In this case the severe collapse of the gorilla population continues to be described by the Eq. 4.

In spite of its simplified well-mixed mass-action kinetics, our results suggest a way on how to minimize the probability of a disastrous collapse in human populations. Humans routinely share pathogens with animals. Indeed, more than half of nearly 1500 known human pathogens are shared with at least one animal host [2]. Wolfe et al. [3] classified such zoonotic diseases into 5 categories (called evolutionary stages) out of which stages 2-4 differ from each other exclusively by their basic reproduction number in human-to-human transmission (*C*_11_ in our notation). Stage 2 is characterized by a complete lack of human-to-human transmission (*C*_11_ = 0), Stage 3 - by sub-critical human-to-human transmission 0 *< C*_11_ *<* 1), and Stage 4 - by super-critical human-to-human transmission (*C*_11_ *>* 1). Human-to-human basic reproduction number, *C*_11_, is clearly important both for endemic state stability considered in Ref. [8] as well as for the epidemic-driven population collapse considered here, especially in the case where the disease does not spread on its own in their animal host (*C*_22_ *<* 1). However, as demonstrated in Fig. 2b, for a pathogen capable to spread in the co-infected animal (*C*_22_ *>* 1 such as example used in Fig. 2b) the human-to-human basic reproduction number *C*_11_ has only mild and qualitative impact on Γ_1_ quantifying the logarithm of the population collapse in humans. Much more important factor is the magnitude of the animal-to-human collapse factor *C*_12_ = *β*_12_*S*_2_(0)*/γ*_2_. It is proportional to the population of the animal host, which could potentially be very large. We are outnumbered by populations of, for example, small birds, rats and mice. Our paper emphasizes the advantage of limiting our exposure to such species with large populations and high growth rates. Perhaps much of the recent trend showing the overall decrease in occurrence of serious epidemics in the industrial world could be attributed to progressively less frequent contacts between humans living in major population centers and these animals. To prevent serious disease outbreaks in the future it may be particularly useful to closely monitor abundant disease carriers in regions with high potential for inter-species contacts.

## ACKNOWLEDGMENTS

We thank Lone Simonsen for inspiring discussions and for bringing to our attention the 2002-2003 Ebola epidemics among gorillas.

